# Synaptic and intrinsic potentiation in O-LM interneurons is induced by theta patterns of stimulation

**DOI:** 10.1101/2022.01.19.476905

**Authors:** Malika Sammari, Yanis Inglebert, Norbert Ankri, Michaël Russier, Salvatore Incontro, Dominique Debanne

## Abstract

Oriens lacunosum-moleculare (O-LM) interneurons display a non-conventional form of long-term synaptic potentiation (LTP) conferred by calcium-permeable AMPA receptors (CP-AMPAR). So far, this form of LTP has been induced in O-LM cells by physiologically unrealistic protocols. We report here the induction of both synaptic and intrinsic potentiation in O-LM interneurons following stimulation of afferent glutamatergic inputs in the theta (θ) frequency range. LTP is induced by synaptic activation of CP-AMPAR whereas long-term potentiation of intrinsic excitability (LTP-IE) results from the mGluR1-dependent down-regulation of Kv7 voltage-dependent potassium channel and hyperpolarization activated and cyclic nucleotide-gated (HCN) channel through the depletion of phosphatidylinositol-4,5-bi-phosphate (PIP2). LTP and LTP-IE are reversible, demonstrating that both synaptic and intrinsic changes are bidirectional in O-LM cells. We conclude that physiological stimuli such as θ patterns induce synaptic and intrinsic potentiation in O-LM interneurons.

## Introduction

Inhibitory interneurons play a fundamental role in brain activity by balancing synaptic and intrinsic excitation and finely orchestrating brain rhythms (Pelkey et al., 2017). This fine tuning achieved by GABAergic interneurons is required for perception and for memory formation (Isaacson and Scanziani, 2011; Kullmann et al., 2012). Hippocampal interneurons represent a diverse population of cell types that inhibit different cellular regions. In the hippocampus, parvalbumin (PV)-expressing interneurons target the perisomatic region of pyramidal cells whereas somatostatin (SOM)-expressing interneurons such as *Oriens Lacunosum Moleculare* (O-LM) cells inhibit the distal dendritic region of pyramidal cells (Klausberger and Somogyi, 2008; Pelkey et al., 2017).

O-LM interneurons are active during theta and gamma oscillations (Klausberger et al., 2003; Pangalos et al., 2013) and participate in feedback inhibitory circuits within area CA1. They receive glutamatergic inputs from CA1 pyramidal cells and in turn, they inhibit distal dendrites of CA1 pyramidal cells and PV interneurons targeting the Schaffer collateral inputs (Leão et al., 2012). Thus O-LM interneurons selectively inhibit information from entorhinal cortex and disinhibit proximal dendrites that receive CA3 inputs (Leão et al., 2012; Müller and Remy, 2014). Thereby, O-LM interneuron activity routes the intra-hippocampal pathway used for the recognition of novelty and extra-hippocampal pathway used for memory retrieval (Andersen, 2006). Thus, any acquired regulation of O-LM activity may favor one or the other pathway.

O-LM interneurons express long-lasting forms of synaptic plasticity. Anti-Hebbian long-term potentiation (LTP) has been reported following high frequency stimulation of CA1 pyramidal axons paired with a post-synaptic hyperpolarization to −90 mV (Lamsa et al., 2007). This form of LTP requires the activation of Ca^2+^-permeable AMPA receptors (CP-AMPAR) that are preferentially open at hyperpolarizing membrane potential and thus function as logical “and not” gates (Kullmann and Lamsa, 2007). In fact, CP-AMPAR are blocked by polyamines during postsynaptic depolarization that occurs during EPSP summation induced by high frequency stimulation (Rozov et al., 1998). However, it is still unclear whether LTP can be induced in O-LM interneurons under more physiological conditions, i.e. at a physiologically plausible membrane potential and using realistic stimulation frequencies.

We show in this study that low frequency stimulations (5 Hz or θ activity) of the axons of CA1 pyramidal cells induce synaptic LTP and long-term potentiation of intrinsic excitability (LTP-IE) in O-LM interneurons. Both LTP and LTP-IE are reversible. LTP is induced by CP-AMPA receptors whereas LTP-IE requires mGluR1 for its induction. The mechanisms underlying expression of LTP-IE involve the down-regulation of both HCN and Kv7 channels through a PLC-dependent depletion of PIP2 and PKC activity. Our results thus demonstrate that physiological stimuli such as θ patterns enhance both synaptic strength and intrinsic excitability in CA1 O-LM interneurons, thus suggesting that *in vivo*, O-LM interneurons may express these two forms of plasticity.

## Results

### Co-induction of LTP and LTP-IE in O-LM cells

As LTP in O-LM cells is classically anti-Hebbian and induced by stimulation of CP-AMPAR, we tested whether low frequency stimulation of glutamatergic inputs induced LTP in these neurons. O-LM interneurons were recorded in the *stratum oriens* of acute slice of P14-21 rat hippocampus with spermine-filled electrodes and were identified by their firing properties (i.e., regular firing, deep AHP, and the presence of a depolarizing sag, **Supplementary Figure 1A**) and their morphology (i.e. their bipolar dendrites emerging from a spindle-like cell body and their long axonal process terminating in the apical region of the CA1 pyramidal cell dendrites; **Figure 1A**). To maximize calcium influx through CP-AMPAR, synaptic stimulation was induced while the neuron was held in current clamp at −77 mV with negative holding current (mean: −24 pA). Synaptic and intrinsic changes were tested using two different protocols. Input/output curves were plotted at the beginning and the end of each experiment and EPSP slope, input resistance (R_in_) and action potential (AP) number were monitored before and after 5 Hz stimulation (**Supplementary Figure 1A**).

**Figure 1.**
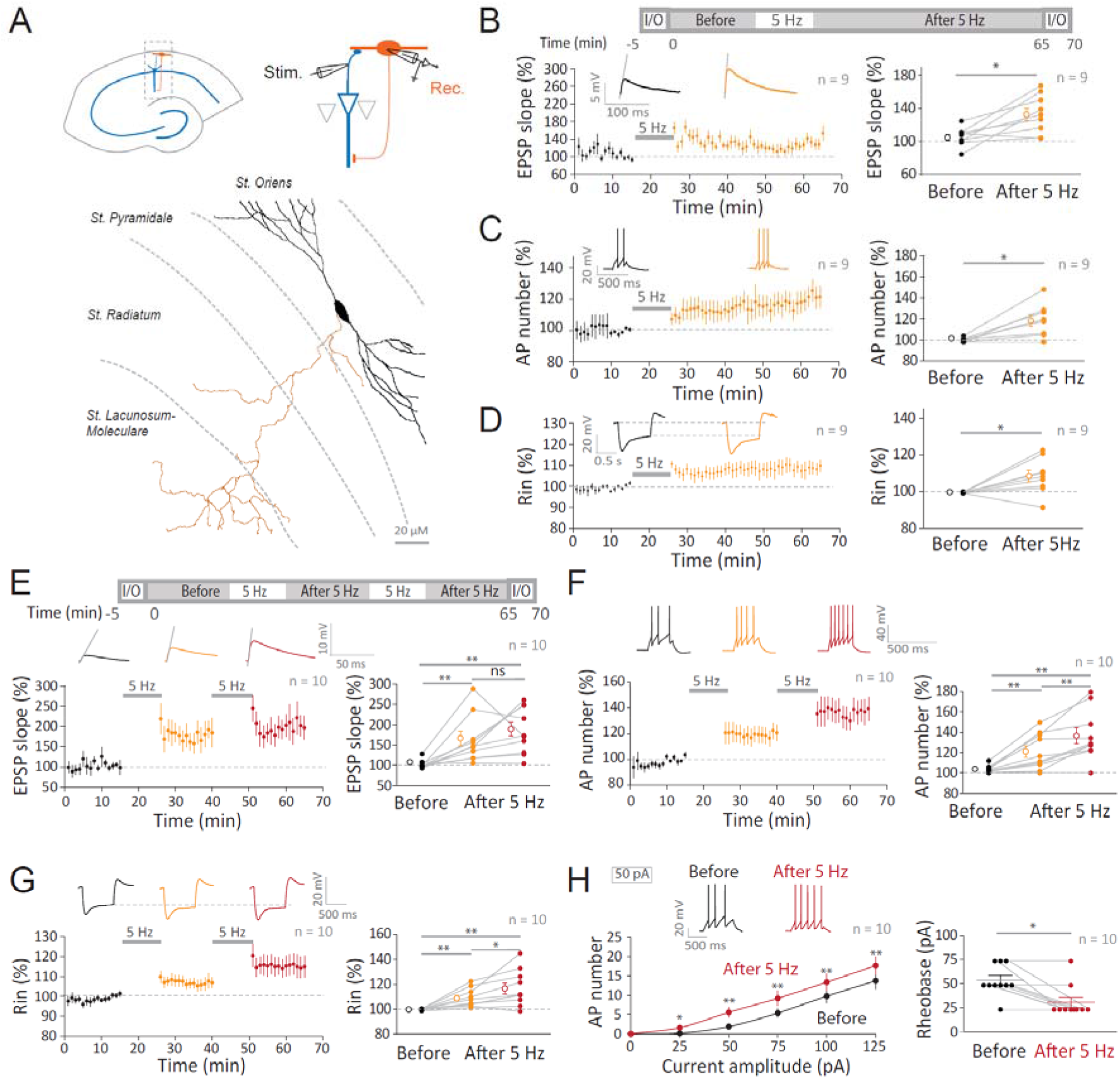
Five Hz stimulation induces LTP and LTP-IE in O-LM interneurons. **A**. Top, recording configuration and morphological identification of O-LM interneuron. Bottom, reconstruction of an O-LM interneuron filled with biocytin. Black segments represent dendrites and orange segments represent axon. **B**. Top, stimulation configuration (I/O, input/output curve; Before and After 5 Hz, test of EPSP slope, R_in_ and AP number, see **Supplementary Figure 1**). Time-course (left) and group data (right) of normalized EPSP slope before and after 5 Hz stimulation. **C**. Time-course (left) and group data (right) of normalized AP number before and after 5 Hz stimulation. **D**. Time-course (left) and group data (right) of normalized R_in_ before and after 5 Hz stimulation. **E**. Top, stimulation configuration using two 5 Hz stimulation episodes. Left, time-course of normalized EPSP slope following 5 Hz stimulation. Right, group data. **F**. Left, time-course of normalized AP number. Right, group data. **G**. Left, time course of normalized R_in_. Right, group data. **H**. Input/output curves (left) and rheobase (right) before and after 2 episodes of 5 Hz stimulation. Wilcoxon and Mann-Whitney tests were used: ns, p>0.05; *, p<0.05; **, p<0.01; ***, p<0.001.

A single episode of 5 Hz stimulation resulted in a long-lasting (>25 min) potentiation of both excitatory synaptic transmission (133 ± 8% of control EPSP slope, n = 9, Wilcoxon test, p < 0.05; **Figure 1B**) and intrinsic excitability (118 ± 5% of control AP number, n = 9, Wilcoxon test, p < 0.05; **Figure 1C**). In addition, R_in_ was found to be elevated (108 ± 3% of the control R_in_, n = 9, Wilcoxon test, p < 0.05; **Figure 1D**). Input/output changes revealed a significant increase in spike number for 50 and 75 pA but not for the other values of current tested (**Supplementary Figure 1B**). In addition, no significant change in the rheobase was observed (**Supplementary Figure 1B**) and the holding current was unchanged (**Supplementary Figure 1C**).

In order to check whether LTP and LTP-IE are already saturated by a single 5 Hz stimulation episode or, on the contrary, able to encode different levels of stimulation, we compared the data after a first stimulation episode to those after a second stimulation. No significant gain in synaptic strength was obtained after the second stimulation episode (162 ± 18% of control EPSP slope after the first stimulation episode and 184 ± 18%, after the second stimulation episode; p > 0.05 Wilcoxon test, **Figure 1E**). In contrast, AP number and R_in_ were found to increase successfully after each episode of stimulation (121 ± 6% and 136 ± 8% of control AP number, **Figure 1F** and 109 ± 2% and 117 ± 5% of control R_in_, **Figure 1G**; p < 0.01 and p < 0.05, Wilcoxon test, n = 10), indicating that, in contrast to LTP, LTP-IE and the associated change in R_in_ are discriminative. Moreover, a significant leftward shift in the input output curve and a reduced rheobase were also found after the two episodes of 5 Hz stimulation (from 55 ± 5 pA before to 33 ± 5 pA after the stimulation, p < 0.05 Wilcoxon test, n=10, **Figure 1H**). The AP threshold was significantly hyperpolarized after the two episodes of 5 Hz stimulation (from −51.9 ± 1.8 mV to −57.4 ± 1.9 mV, n = 10; p < 0.01; **Supplementary Figure 2A**). In addition, input resistance tested with depolarizing subthreshold pulses of current was found to increase after 5 Hz stimulation (**Supplementary Figure 2B**). Interestingly, stimulation at 10 Hz induced LTP (123 ± 7% and 132 ± 11%, n = 9) but not LTP-IE (98 ± 2% and 98 ± 3%, n = 9; **Supplementary Figure 3**), suggesting that induction of LTP and LTP-IE might involve separate pathways. We conclude that low frequency stimulation of glutamatergic inputs at 5 Hz induces both synaptic and intrinsic potentiation in O-LM interneurons maintained near their resting membrane potential.

### Bidirectional synaptic and intrinsic plasticity in O-LM cells

Synaptic transmission and intrinsic excitability are depressed following a negative pairing (NP) protocol delivered at 10 Hz and consisting in pairing EPSP with single postsynaptic AP with a delay of −10 ms (Incontro et al., 2021). We therefore tested whether synaptic and intrinsic changes induced by 5 Hz stimulation were reversible. LTP and LTP-IE were first induced by a single episode of 5 Hz stimulation (145 ± 16% of the control EPSP slope, n = 7, Wilcoxon test p < 0.05, and 113 ± 5% of the control AP number, n = 7, Wilcoxon test p < 0.05; **Figure 2A** and **2B**). Then, NP at 10 Hz was applied as previously shown (Incontro et al., 2021; Péterfi et al., 2012). Importantly, both LTP and LTP-IE were reversed by induction of synaptic and intrinsic depotentiation (**Figure 2A** and **2B**). In addition, a significant shift in the input-output curve was observed associated with a non-significant bidirectional change in the rheobase (**Figure 2C**). We conclude that synaptic and intrinsic plasticity are reversible in O-LM interneurons.

**Figure 2.**
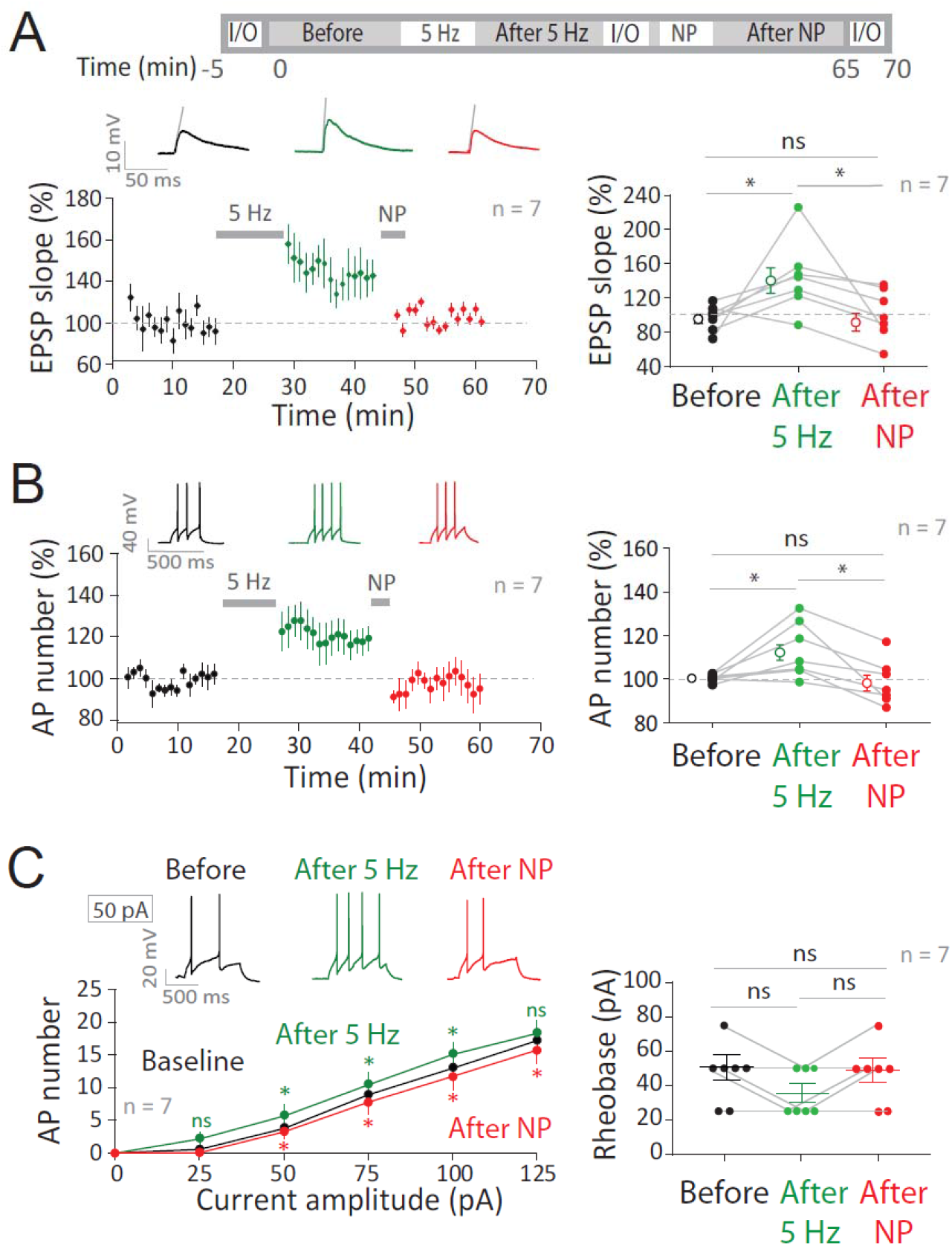
Bidirectional synaptic and intrinsic plasticity in O-LM interneurons. **A**. Top, protocol of stimulation. Bottom left, time-course of the normalized EPSP slope showing LTP induced by 5 Hz and depotentiation induced by negative pairing (NP). Right, group data. **B**. Left, time-course of the normalized AP number. Right, group data. **C**. Input/output curves (left) and rheobase (right) before and after 5 Hz and after NP. Statistics related before vs. after 5 Hz data are in green and those related to after 5 Hz vs. after NP data are in red. Wilcoxon test was used: ns, p>0.05; *, p<0.05.

### Theta activity induces LTP and LTP-IE in O-LM interneurons

Low frequency stimulations of pyramidal cells induce LTP and LTP-IE in O-LM interneurons. However, it is still unknown if physiological activity of pyramidal cells as theta (θ) activity could lead to synaptic and intrinsic plasticity in O-LM interneurons. To determine the pattern of pyramidal cell discharge during θ waves, pyramidal cell activity was recorded in patch-clamp under physiological conditions (external solution contained 1.3 mM Ca^2+^ and 1 mM Mg^2+^(Inglebert et al., 2020)) and in the presence of 25 μM carbachol (**Figure 3A**). During θ oscillation, the discharge pattern had a mean frequency of ~5 Hz. The recorded θ pattern was then used to stimulate pyramidal cell axon while an O-LM cell was recorded (**Figure 3B**). Interestingly, after stimulation of pyramidal cell axons using 2 episodes of θ-like pattern in 3 mM Ca^2+^ and 2 mM Mg^2+^, O-LM interneurons expressed LTP (144 ± 15% and 169 ± 24% of EPSP slope control, p < 0.01 and p < 0.05 Wilcoxon test, n = 8, **Figure 3C**) and LTP-IE (104 ± 2% and 113 ± 6% of AP number control, p < 0.05 Wilcoxon test, n = 8, **Figure 3D**). Furthermore, an increase of R_in_ (106 ± 2% and 108 ± 2% of R_in_ control, p < 0.05 Wilcoxon test, n = 8, **Figure 3E**) with a significant leftward shift in the input/output curve (**Figure 3F**) was observed. We conclude that stimulation of pyramidal cells at θ frequency induces synaptic and intrinsic potentiation in O-LM interneurons.

**Figure 3.**
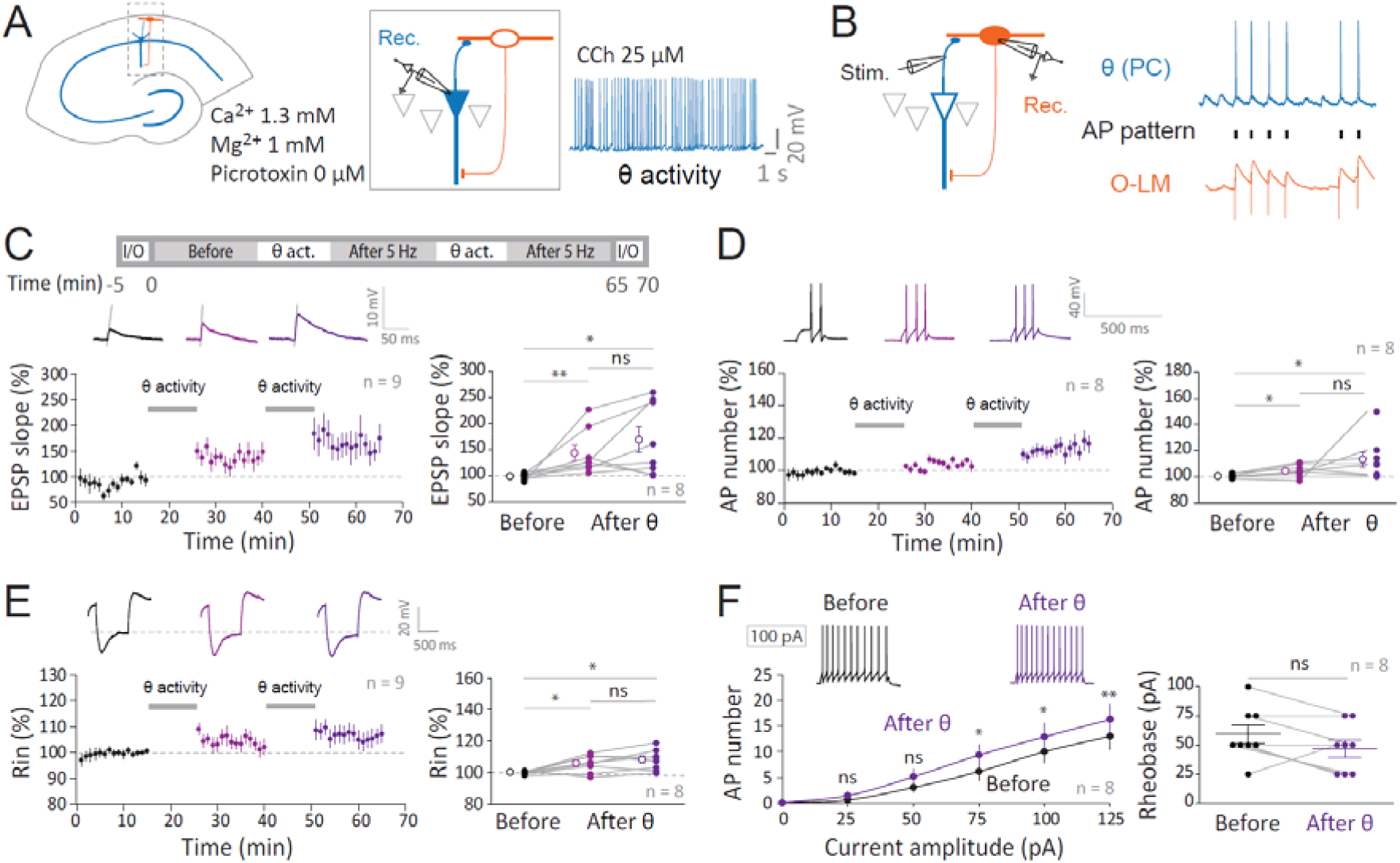
θ stimulation induces LTP and LTP-IE in O-LM interneurons. **A**. Recording configuration and discharge pattern of CA1 pyramidal cells during θ oscillations induced by carbachol (CCh) in physiological external Ca^2+^ and Mg^2+^ concentration. **B**. Stimulation of pyramidal cell axons with the θ pattern while O-LM interneuron was recorded. **C**. Top, experimental protocol. Bottom, time-course (left) and group data (right) of normalized EPSP slope before and after θ stimulation. **D**. Time-course (left) and group data (right) of normalized AP number before and after θ stimulation. **E**. Time-course (left) and group data (right) of normalized R_in_ before and after θ stimulation. **F**. Input/output curve (right) and rheobase (left) before and after θ stimulation. Wilcoxon and Mann-Whitney tests were used: ns, p>0.05; *, p<0.05; **, p<0.01.

### CP-AMPA receptor is required for LTP induction and mGluR1 for LTP-IE induction

Next, we tested the induction mechanisms of both LTP and LTP-IE in O-LM interneurons. We first examined whether LTP induced by 5 Hz stimulation required the involvement of CP-AMPAR using the selective antagonist of CP-AMPAR, NASPM (100 μM). In the presence of NASPM, LTP was abolished (102 ± 3% of the control EPSP after the first stimulation episode and 92 ± 4%, n = 10 after the second stimulation episode, **Figure 4A**). Indeed, compared to the control situation, the synaptic potentiation was significantly reduced in NASPM (p<0.01 and 0.001, **Figure 4A**). However, LTP-IE was still present (131 ± 11% and 132 ± 11%, n = 10) and thus reached a level comparable to the control (p > 0.1; **Figure 4B**). Moreover, as in control, the input-output curve in the presence of NASPM was shifted leftward but the rheobase was not significantly reduced (45 ± 5 pA vs. 37 ± 4 pA after the second stimulation, n = 10, Wilcoxon, p > 0.1; **Figure 4C**). We conclude that LTP but not LTP-IE depends on synaptic activation of CP-AMPAR.

**Figure 4.**
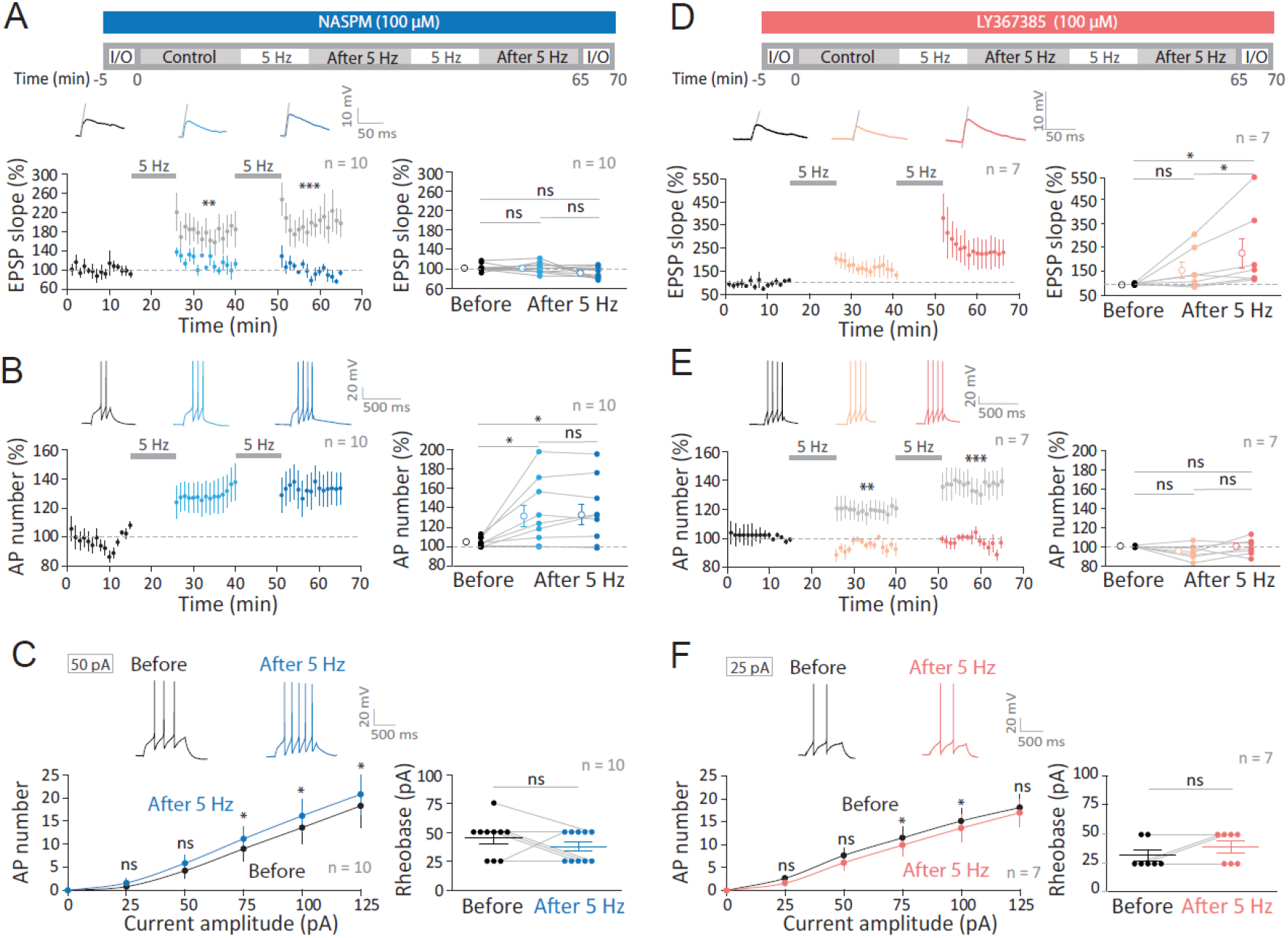
LTP induction requires CP-AMPAR whereas LTP-IE induction requires mGluR1. **A**. Top, experimental protocol in the presence of the CP-AMPA receptor antagonist, NASPM (100 μM). Bottom, time-course (left) and group data (right) of the normalized EPSP slope before and after 5 Hz stimulation in the presence of NASPM. The grey data correspond to the normalized EPSP slope after 5 Hz stimulation obtained in control. Note the significant difference. **B**. Time-course (left) and group (right) of normalized AP number before and after 5 Hz stimulation in the presence of NASPM. **C**. Input/output curve (right) and rheobase (left) before and after 5 Hz stimulation in presence of NASPM. **D**. Top, experimental protocol in the presence of the mGluR1 receptor antagonist, LY367385 (100 μM). Bottom, time-course (left) and group data (right) of the normalized EPSP slope before and after 5 Hz stimulation in the presence of LY-367385. **E**. Time-course (left) and group data (right) of normalized AP number before after 5 Hz stimulation in the presence of LY-367385. The grey data correspond to the normalized AP number after 5 Hz stimulation obtained in control. Note the significant difference. **F**. Input/output curves (right) and rheobase (left) before and after 5Hz stimulation in presence of LY367385. Wilcoxon and Mann-Whitney tests were used: ns, p>0.05; *, p<0.05; **, p<0.01; ***, p<0.001.

Metabotropic glutamate receptors type 1 (mGluR1) are also required for induction of LTP in O-LM interneurons induced by high frequency presynaptic stimulation paired with postsynaptic hyperpolarization (Le Duigou and Kullmann, 2011; Perez et al., 2001). We therefore tested whether LTP induced by 5 Hz stimulation also depends on mGluR1. In the presence of the mGluR1 inhibitor LY367385 (100 μM), LTP was induced (158 ± 32%, n = 7 and 228 ± 62%, n = 7; p < 0.05; **Figure 4D**) to a level comparable to controls (p > 0.1). However, LTP-IE was totally blocked in the presence of LY367385 (95 ± 3%, n = 7 and 100 ± 3%, n = 7; p > 0.1; **Figure 4E**) and was significantly different from the control LTP-IE (p < 0.01 and p < 0.001). Furthermore, a slight rightward shift in the input-output curve associated with a non-significant increase in the rheobase was observed (32 ± 5 pA and 39 ± 5 pA, p > 0.1; **Figure 4F**). We conclude that CP-AMPAR is required for LTP induction and mGluR1 for LTP-IE.

### Down-regulation of HCN and Kv7 channel accounts for LTP-IE in O-LM cells

O-LM interneurons express HCN and Kv7 channels (Gastrein et al., 2011; Lawrence et al., 2006). HCN channels are regulated in long-term changes in excitability (Brager and Johnston, 2007; Campanac et al., 2008; Fan et al., 2005; Gasselin et al., 2015, 2017). In addition, Kv7 channels are involved in the reduction of intrinsic excitability observed after LTD induction (Incontro et al., 2021). We therefore measured both currents in voltage-clamp before and after 2 episodes of 5 Hz stimulation. A significant reduction of both the HCN and Kv7 channel-mediated currents was observed following stimulation (**Figure 5A**). To confirm the implications of both channels, we used specific blockers of each channels. In the presence of 1 μM ZD-7288, a specific blocker of HCN channels, LTP comparable to control (p > 0.1) was still induced (150 ± 10% and 145 ± 8%, n = 9; **Supplementary Figure 4A**). In contrast, LTP-IE was partially occluded (106 ± 5% and 115 ± 5%; n = 9, **Figure 5B**), suggesting the contribution of HCN channels in the expression of LTP-IE. A small but significant leftward shift in the input-output curve was observed (**Figure 5C**), suggesting that another ion channel is involved in the change in excitability. In the presence of 10 μM XE-991, a specific blocker of Kv7 channels, LTP was still induced (138 ± 14% and 158 ± 23%, n = 8; **Supplementary Figure 4B**) but LTP-IE was also partially occluded (107 ± 3% and 113 ± 4%, n = 8; **Figure 5D**). Similarly, a small but significant shift in the input-output curve was observed (**Figure 5E**). Interestingly, R_in_ increased in the presence of XE-991 (108 ± 3% and 118% ± 3%) but not in the presence of ZD-7288 (98 ± 2% and 99 ± 3%). Next, we repeated the experiment in the presence of both ZD-7288 and XE-991. No change in LTP was observed (146 ± 11% and 155 ± 10%, n = 10; **Supplementary Figure 4C**) but LTP-IE was absent (101 ± 4% and 107 ± 4%, n = 10; **Figure 5F**) and input-output curves were indistinguishable (**Figure 5G**). Altogether, these results indicate that LTP-IE is expressed through the down-regulation of both HCN and Kv7 channels.

**Figure 5.**
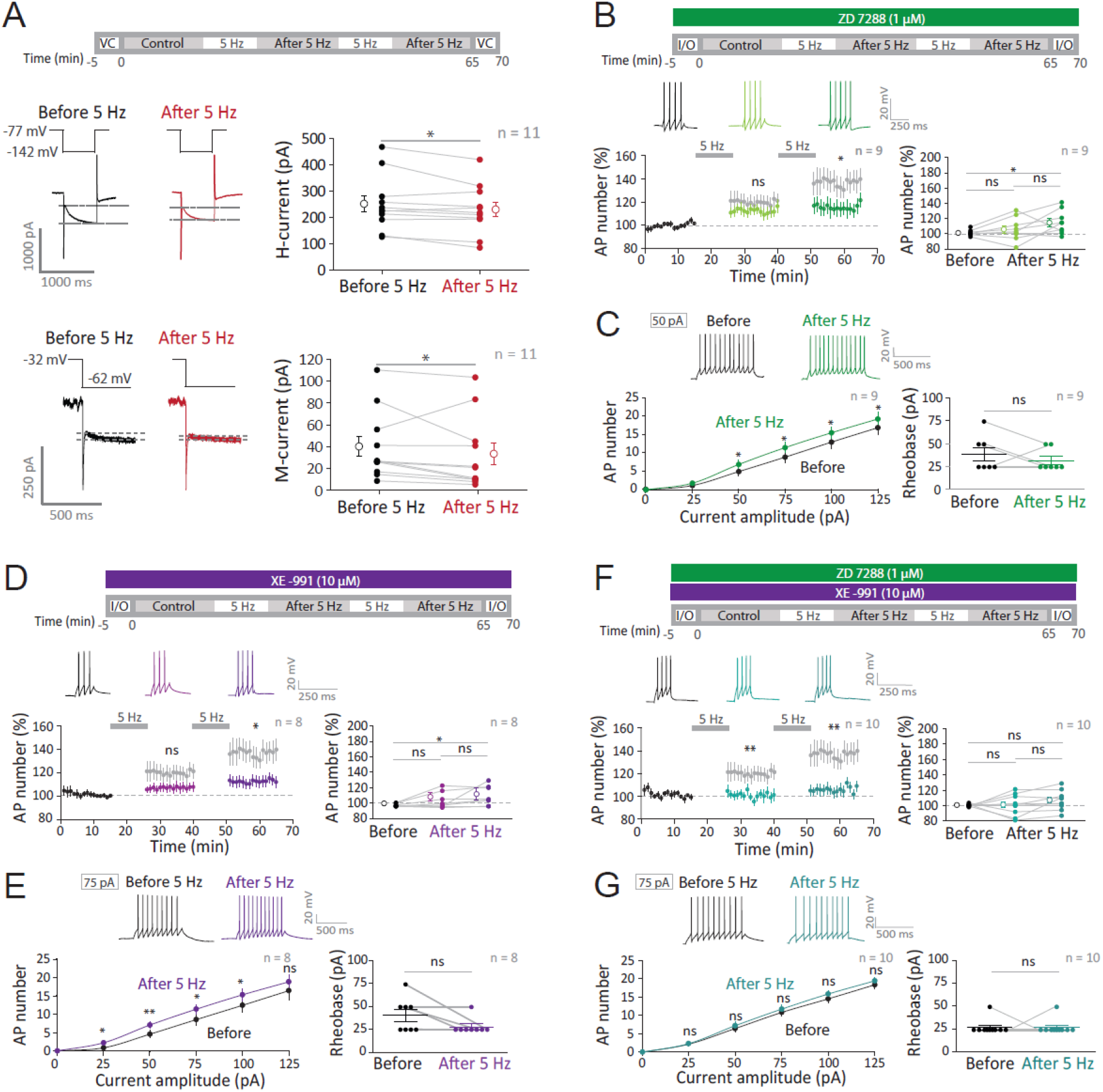
Expression of LTP-IE: down-regulation of HCN and Kv7 channels. **A**. Voltage-clamp measurements of HCN and Kv7 channel-mediated currents before and after 5 Hz. Top, experimental protocol. Middle, reduction of the HCN channel-mediated current evoked by a hyperpolarizing current step from −77 mV to −142 mV. Left, representative traces. Right, group data. Bottom, reduction of the Kv7 channel-mediated M-type current evoked by a hyperpolarization from −32 mV to −62 mV. Left, representative traces. Right, group data. **B**. Top, experimental protocol in the presence of the HCN channel inhibitor, ZD-7288 (1 μM). Bottom, time-course (left) and group data (right) of normalized AP number before and after 5 Hz stimulation in the presence of ZD-7288. The grey data correspond to the normalized AP number after 5 Hz stimulation obtained in control. Note the significant difference. **C**. Input/output curves (left) and rheobase (right) before and after 5 Hz stimulation in the presence of ZD-7288. **D**. Top, experimental protocol in the presence of the Kv7 channel inhibitor, XE-991 (10 μM). Bottom, time-course (left) and group data (right) of normalized AP number before and after 5 Hz stimulation in the presence of XE-991. The grey data correspond to the normalized AP number after 5 Hz stimulation obtained in control. **E**. Input/output curve (right) and rheobase (left) before and after 5 Hz stimulation in the presence of XE-991. Time-course (left) and group data (right) of normalized AP number before and after 5 Hz stimulation in the presence of XE-991. **F**. Top, experimental protocol in the presence of ZD-7288 and XE-991. Bottom, time-course (left) and group data (right) of normalized AP number before and after 5 Hz stimulation in the presence of ZD-7288 and XE-991. The grey data correspond to the normalized AP number after 5 Hz stimulation obtained in control. **G**. Input/output curves (right) and rheobase (left) before and after 5 Hz stimulation in the presence ZD-7288 and XE-991. Wilcoxon and Mann-Whitney tests were used: ns, p>0.05; *, p<0.05; **, p<0.01; ***, p<0.001.

### LTP-IE involves PIP2 depletion through the activation of PLC

Both Kv7 and HCN channels need phosphatidylinositol-4,5-biphosphate (PIP2) to be active (Zhang et al., 2003; Zolles et al., 2006). Thus, a possible scenario could be that stimulation of mGluR1 activates the phospholipase C (PLC), leading to the depletion of PIP2 and thus to the reduction of the Kv7-mediated M-current and the HCN-mediated H-current. We first tested the role of PLC in both LTP and LTP-IE induction. In the presence of 10 μM U-73122, a specific inhibitor of PLC, LTP reached normal value (131 ± 7% and 141 ± 12%, n = 8; **Figure 6A**), but LTP-IE was blocked (103 ± 2% and 106 ± 3%, n = 8; **Figure 6B**) and input-output curves were unchanged (**Figure 6D**). Interestingly, R_in_ was not significantly elevated (104 ± 2% and 103 ± 3%, n = 8; **Figure 6C**), indicating that in addition to Kv7 channel, HCN channel is also targeted by PLC.

**Figure 6.**
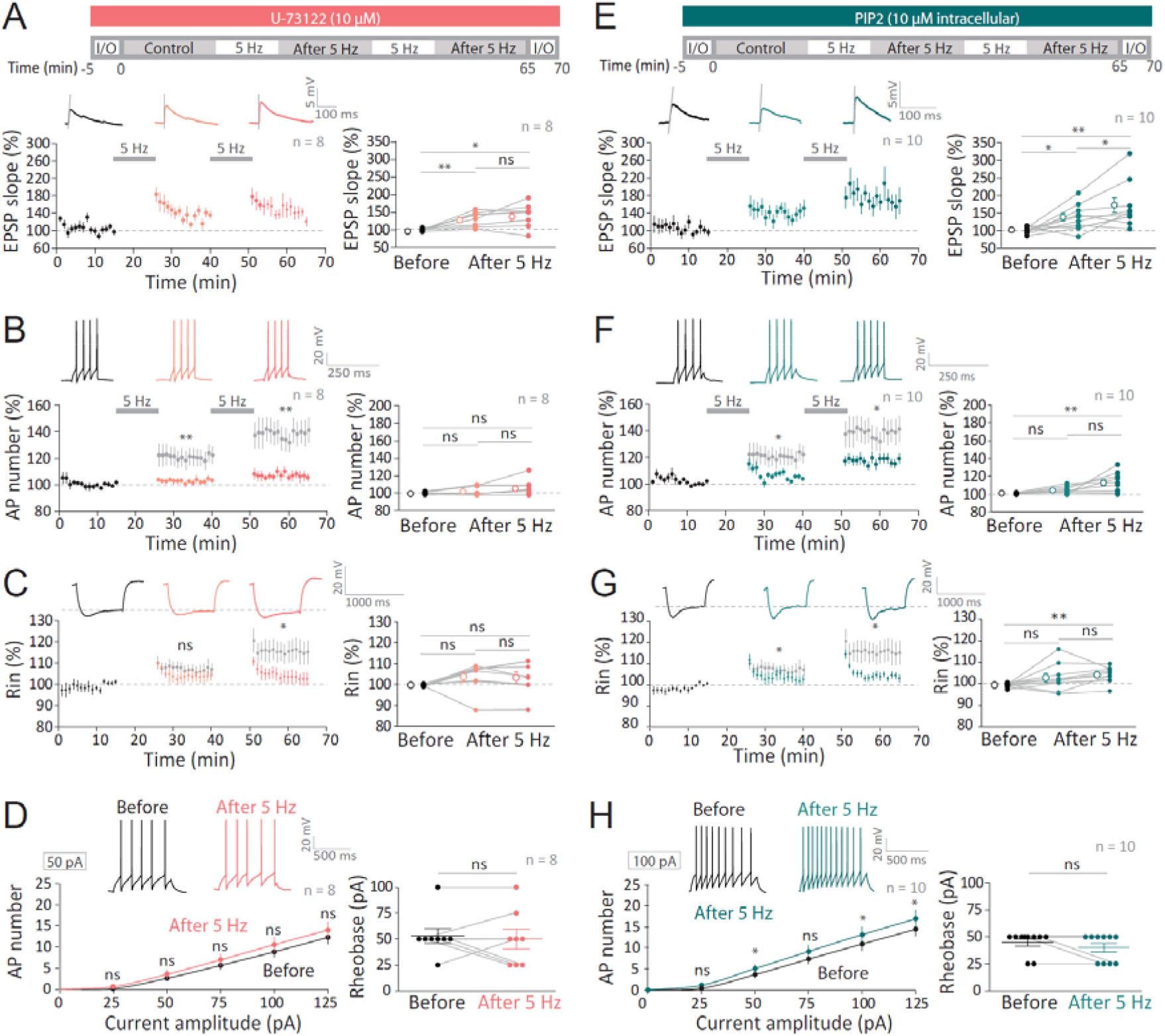
LTP-IE requires PLC activation and PIP2. **A**. Top, experimental protocol in the presence of the PLC inhibitor, U-73122 (10 μM). Bottom, time-course (left) and group data (right) of normalized EPSP slope before and after 5 Hz stimulation in the presence of U-73122. **B**. Time-course (left) and group data (right) of normalized AP number before and after 5 Hz stimulation in the presence of U-73122. Grey data correspond to the normalized AP number after 5 Hz stimulation obtained in control. Note the significant difference. **C**. Time-course (left) and group data (right) of normalized R_in_ before and after 5 Hz stimulation in the presence of U-73122. Grey data correspond to the normalized R_in_ after 5 Hz stimulation obtained in control. Note the significant difference. **D**. Input/output curves (right) and rheobase (left) before and after 5 Hz stimulation in the presence of U-73122. **E**. Top, experimental protocol with PIP2 (10 μM) in the recording pipette. Bottom, time-course (left) and group data (right) of normalized EPSP slope before and after 5 Hz stimulation with PIP2 in the recording pipette. **F**. Time-course (left) and group data (right) of normalized AP number before and after 5 Hz stimulation with PIP2 in the recording pipette. Grey data correspond to the normalized AP number after 5 Hz stimulation obtained in control. Note the significant difference. **G**. Time-course (left) and group data (right) of normalized R_in_ before and after 5 Hz stimulation with PIP2 in the recording pipette. Grey data correspond to the normalized R_in_ after 5 Hz stimulation obtained in control. Note the significant difference. **H**. Input/output curve (right) and rheobase (left) before and after 5 Hz stimulation with PIP2 in the recording pipette. Wilcoxon and Mann-Whitney tests were used: ns, p>0.05; *, p<0.05; **, p<0.01.

Next, PIP2 was added to the recording pipette in order to reduce the depletion of PIP2 on Kv7 and HCN channels. In these conditions, LTP was comparable to the control (137 ± 12% and 170 ± 20%, n = 10; **Figure 6E**), but interestingly, LTP-IE was reduced (104 ± 1% and 113 ± 3%, n = 10; **Figure 6F**) and the input-output curve was weakly shifted (**Figure 6H**), indicating that PIP2 supply prevents the regulation of Kv7 and HCN channels. Interestingly, here again R_in_ was not significantly increased (103 ± 2% and 104 ± 1%; **Figure 6G**), indicating that HCN channels are sensitive to the depletion of PIP2. Altogether, these results indicate that LTP-IE involves the depletion of PIP2 through the activation of PLC to reduce the currents mediated by both HCN and Kv7 channels.

### Inhibition of PKC prevents both LTP and R_in_ increase

HCN currents are also inhibited by activation of PKC (Williams et al., 2015). PKC can be stimulated by Ca^2+^. Furthermore, in addition to the PIP2 depletion, activation of PLC may also activate PKC through the mobilization of Ca^2+^ by IP3. We therefore tested the effect of PKC inhibition on both LTP and LTP-IE induced by 5 Hz stimulation. In the presence of 10 μM chelerythrine, a cell permeable inhibitor of PKC, LTP was strongly reduced (131 ± 17% and 117 ± 16%, n = 9; **Figure 7A**), and LTP-IE was reduced compared to control (111 ± 3% and 117 ± 3%, n = 9; **Figure 7B**). Importantly, R_in_ was unchanged (101 ± 2% and 102 ± 4%, n = 9; **Figure 7C**), suggesting that the regulation of HCN channels is frozen in the presence of the PKC inhibitor. However, the input-output curve was significantly shifted leftward (**Figure 7D**). We conclude that PKC is involved in both LTP and LTP-IE through the regulation of HCN channels.

**Figure 7.**
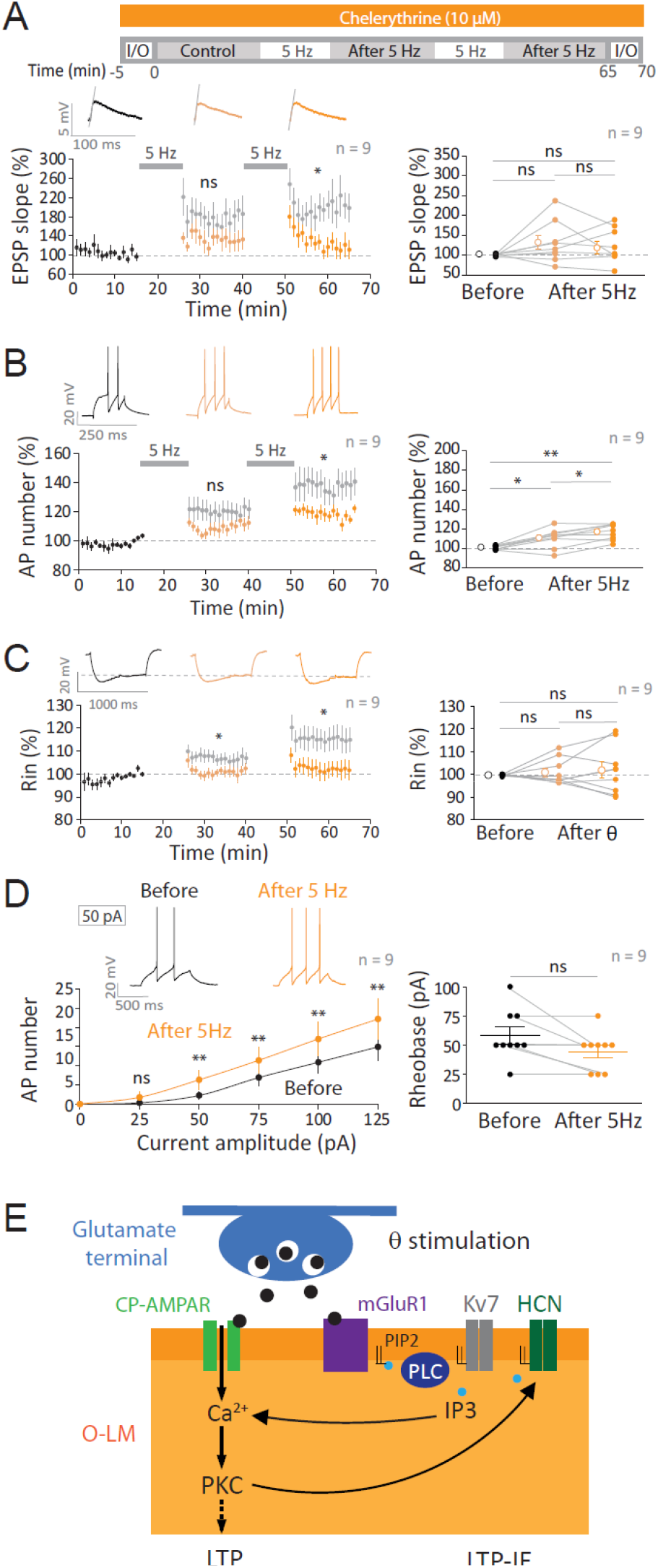
PKC inhibition blocks LTP and reduces LTP-IE through HCN channels. **A**. Top, stimulation protocol in the presence of the PKC inhibitor, chelerythrine (10 μM). Left, time-course of the normalized EPSP slope. Note the significant reduction compared to control (grey data points). Right, group data. **B**. Left, time-course of the normalized AP number. Note the significant reduction compared to control (grey data). Right, group data. **C**. Left, time-course of the normalized R_in_. Note the significant reduction compared to control (grey data). Right, group data. **D**. Input/output curves (left) and rheobase (right) in the presence of chelerythrine. **E**. Summary scheme of the induction and expression mechanisms of LTP and LTP-IE by 5 Hz stimulation in O-LM interneurons. A glutamatergic input is represented in blue and the dendrites of O-LM cell in orange. Release of glutamate (black dots) activate CP-AMPAR and mGluR1. CP-AMPAR is permeable to Ca^2+^ ions that activate PKC and lead to LTP induction. mGluR1 stimulation activate PLC that depletes PIP2. PIP2 depletion closes both Kv7 and HCN channels. Wilcoxon and Mann-Whitney tests were used: ns, p>0.05; *, p<0.05; **, p<0.01.

## Discussion

We show here that low frequency stimulation at 5 Hz (or θ stimulation pattern) induces both LTP and LTP-IE in rat O-LM interneurons. These two modifications are synergistic and provide a high level of functional redundancy. Our study indicates that mechanistically, synaptic and intrinsic potentiation in O-LM interneurons depend on distinct classes of glutamate receptors. LTP induction requires activation of GluA2 lacking CP-AMPAR whereas LTP-IE induction requires the synaptic stimulation of mGluR1 (**Figure 7E**). LTP-IE is expressed through the down-regulation of both HCN and Kv7 channels by the PLC-dependent depletion of PIP2 (**Figure 7E**). In addition, LTP and HCN regulation depend on PKC. LTP and LTP-IE were found to be fully reversible by a protocol that induces LTD and LTD-IE, demonstrating that both synaptic and intrinsic changes are bidirectional. Modulation of intrinsic excitability can potentially have very important consequences on the local hippocampal circuit by modulating the strength of the feedback loop O-LM interneurons establish with CA1 pyramidal neurons.

### θ stimulation patterns induce synaptic and intrinsic plasticity in O-LM cells

We show here that 5 Hz stimulations or θ stimulation patterns during 10 minutes induce both LTP and LTP-IE in O-LM interneurons. θ stimulation patterns were recorded in CA1 pyramidal neurons *in vitro* and used to stimulate excitatory inputs of O-LM interneurons. Each θ episode lasted ~1 minute *in vitro*, but induction of LTP and LTP-IE required 10 minutes of stimulation. Our protocol is nevertheless totally plausible as θ episodes may last several minutes *in vivo* (Malezieux et al., 2020). Stimulation of the Schaffer collaterals at θ frequency induces LTD in pyramidal CA1 neurons (Dudek and Bear, 1992; Mulkey and Malenka, 1992). Furthermore, LTP or LTD can be induced during θ oscillation elicited by stimulation of cholinergic receptors (Huerta and Lisman, 1993, 1995).

### Synergy between synaptic and intrinsic plasticity in O-LM cells

Synaptic and intrinsic plasticity are generally synergistic in hippocampal pyramidal neurons (Campanac and Debanne, 2008; Daoudal et al., 2002; Debanne et al., 2019; Wang et al., 2003). Do GABAergic interneurons also share this rule? Regarding parvalbumin positive basket cells, LTP and LTP-IE are induced conjointly (Campanac et al., 2013), thus reinforcing the synaptic potentiation. O-LM interneurons also support this rule as both LTP and LTD are associated with synergistic changes in excitability. In further support, synaptic and intrinsic depression are both induced by negatively pairing postsynaptic firing with EPSPs (Incontro et al., 2021). Here, we show that the counterpart is also true in O-LM cells as synaptic and intrinsic potentiation are induced by low frequency stimulation such as 5 Hz or θ stimulation pattern. This consideration has been recently extended to homeostatic plasticity by showing that homeostatic regulation of both excitability and synaptic transmission is mediated by the regulation of Kv1 channels in the axon initial segment and at the terminal (Zbili et al., 2021).

### LTP in O-LM interneurons

So far, anti-Hebbian LTP was induced in O-LM interneurons by pairing synaptic stimulation at high frequency (100 Hz) or low frequency (5 Hz) with a post-synaptic hyperpolarization at −90 mV (Lamsa et al., 2007; Le Duigou and Kullmann, 2011; Le Duigou et al., 2015; Nicholson and Kullmann, 2021; Oren et al., 2009). As a membrane potential of −90 mV is hardly obtained in natural conditions, these induction protocols cannot be considered as physiological. We show here that 5 Hz stimulation at a membrane potential of −77 mV is sufficient to induce LTP. mGluR1 and CP-AMPAR are the two main glutamate receptors involved in LTP induction in O-LM interneurons (Lamsa et al., 2007; Le Duigou and Kullmann, 2011) and in perisomatic inhibitory interneurons (Hainmueller et al., 2014). The sensitivity to membrane potential is conferred by CP-AMPAR because of the polyamine-mediated rectification that favors calcium entry at hyperpolarized potentials. Our data reveal that LTP induced by 5 Hz stimulation requires CP-AMPAR as it was blocked by NASPM. However, mGluR1 is not required for LTP induced by 5 Hz since in the presence of LY367385, LTP was similar to control. CP-AMPAR activation induces a calcium influx that is proposed to activate PKC. Interestingly, PKC inhibition significantly reduced LTP magnitude.

Calcium influx is also mediated by nicotinic acetylcholine receptors and especially by α7 nicotinic acetylcholine (α7nACh) receptor (Shen and Yakel, 2009). Interestingly, anti-Hebbian LTP in O-LM cells also requires α7nACh receptor for its induction (Griguoli et al., 2013). Thus, activation of PKC by calcium may be also mediated by α7nACh receptor.

### Intrinsic plasticity in GABAergic interneurons

Activity-dependent changes in intrinsic excitability have been reported in only a few type of hippocampal interneurons including basket cells (Campanac et al., 2013), GABAergic interneuron-targeting neurogliaform interneurons (Chittajallu et al., 2020) and O-LM interneurons (Incontro et al., 2021). Intrinsic plasticity is induced by glutamatergic synaptic stimulation or electrical stimulation and requires the regulation of inhibitory ion channels. LTP-IE in parvalbumin positive basket cells is mediated by synaptic stimulation of mGluR5 induced by high frequency stimulation of glutamatergic inputs and the down-regulation of Kv1 channels (Campanac et al., 2013). Long-lasting increase in excitability of neurogliaform interneurons is induced by action potential firing at 30 Hz and mediated by the regulation of Kv4 channels (Chittajallu et al., 2020). LTD-IE in O-LM cells is induced by negative pairing of single spike with EPSP at 10 Hz and requires the eCB-mediated up-regulation of Kv7 channels (Incontro et al., 2021). We show here that LTP-IE in O-LM interneurons is induced by mGluR1 stimulation and is expressed through the down-regulation of both HCN and Kv7 channels.

Three lines of evidence indicate that HCN channels is involved in LTP-IE. A significant increase in R_in_ was observed following 5 Hz or θ stimulation, the hyperpolarization-activated HCN-current was decreased after 5 Hz stimulation and LTP-IE was reduced in the presence of ZD-7288. Regarding the contribution of Kv7 channels, we show that the voltage-deflection induced with subthreshold positive pulses of current was increased. In addition, the relaxation of the M-current was reduced and LTP-IE was reduced in the presence of XE-991. Importantly, when the two channel inhibitors were co-applied, LTP-IE was prevented, further indicating the involvement of both HCN and Kv7 channels in LTP-IE expression. The lack of change in holding current is also compatible with the regulation of both channels as down-regulation of HCN and Kv7 channels has opposite effects on the resting membrane potential. The hyperpolarization of the AP threshold is also compatible with a reduction in Kv7 channels as the pharmacological blockade of these channels is known to hyperpolarize the AP threshold (Mateos-Aparicio et al., 2014).

O-LM interneurons express both Kv7.2/3 (Lawrence et al., 2006) and HCN2 in their dendrites (Hilscher et al., 2019). Interestingly, mice lacking HCN2 in O-LM interneurons express a higher level of LTP induced at temporoammonic inputs from the entorhinal cortex impinging on distal dendrites of CA1 pyramidal neurons (Matt et al., 2011), suggesting that intrinsic plasticity in O-LM interneurons may lead to the modulation of LTP at distal synapses.

### Lipid-dependent regulation of intrinsic excitability

As proteins inserted in a lipid membrane, ion channels are likely to be highly regulated by lipids. We show here that the down-regulation of both HCN and Kv7 channels involve a PLC-dependent depletion of PIP2. Inhibition of the PLC prevented LTP-IE and R_in_ increase in O-LM interneurons. In addition, PIP2 supply in the recording pipette also suppressed the R_in_ increase and reduced LTP-IE. PIP2 is necessary for the Kv7 channel function (Zhang et al., 2003) and facilitates opening of HCN channels (Zolles et al., 2006). Thus, our findings provide another example of the role of lipid regulation of voltage-gate ion channel in the context of long-lasting plasticity. In fact, we previously showed that eCB such as 2-AG was able to directly bind to Kv7 channel to facilitate its opening (Incontro et al., 2021).

### Functional implications

The fact that excitability of O-LM interneurons can be finely tuned through the reduction of both HCN and Kv7 channels may have important implications at the level of the O-LM cell and for the hippocampal circuit. First, at the cellular level, down-regulation of both HCN and Kv7 channels may decrease the intrinsic resonance in O-LM cells in the depolarizing and hyperpolarizing range. In fact, both Kv7 and HCN channels are responsible for intrinsic resonance at two specific voltage levels (Hu et al., 2002) and O-LM cells display intrinsic resonance (Gastrein et al., 2011). At the local circuit level, enhanced excitability of O-LM interneurons may increase inhibition at distal dendrites of CA1 pyramidal neurons and thus reduce LTP at temporoammonic pathway (Leão et al., 2012). In parallel, the increased inhibition of *stratum radiatum* interneurons may consequently facilitate LTP induction at Schaffer collateral pathway (Leão et al., 2012). In addition, potentiation of the inhibitory synapse established by O-LM with the CA1 pyramidal neuron decreases spiking in response to stimulation of temporoammonic inputs (Udakis et al., 2020).

In the neocortex, the equivalent of O-LM interneurons are Martinotti cells as both cell types express somatostatin and target apical dendrites of principal neurons. Martinotti cells express CP-AMPAR (Lalanne et al., 2016), mGluR1 (Maksymetz et al., 2021) and HCN channels (Wang et al., 2004; Williams and Hablitz, 2015). Therefore, there are good reasons to believe that LTP and LTP-IE might be induced in Martinotti cells. Future studies will certainly provide answers to all these questions.

## Methods

### Acute slices of rat hippocampus

All experimental procedures followed institutional guidelines for the care and use of laboratory animals (Council Directive 86/609/EEC and French National Research Council) and were approved by the local health authority (Préfecture des Bouches-du-Rhône, Marseille). Briefly, 14- to 21-day-old Wistar rats (Charles River) of either sex were anesthetized with isoflurane and decapitated. Hippocampal slices (350 μm) were cut in a N-methyl-D-glucamine (NMDG)-solution containing (in mM: 92 NMDG, 2.5 KCl, 1.2 NaH_2_PO_4_, 30 NaHCO_3_, 20 HEPES, 25 Glucose, 5 Sodium Ascorbate, 2 Thiourea, 3 Sodium Pyruvate, 10 MgCl_2_, 0.5 CaCl_2_) on a vibratome (Leica VT1200S). They were transferred in an NMDG-solution for 10 min maintained at 37°C before resting for 1h at room temperature in oxygenated (95% O_2_/5% CO_2_) artificial cerebro-spinal fluid, ACSF (in mM: 125 NaCl, 2.5 KCl, 0.8 NaH_2_PO_4_, 26 NaHCO_3_, 3 CaCl_2_, 2 MgCl_2_ et 10 Glucose).

### Electrophysiology and data acquisition

Because synaptic inhibition was blocked by picrotoxin (PiTX, 100 μM), the CA3 area was removed surgically to avoid epileptiform activity. Each slice was transferred to a temperature-controlled (30°C) recording chamber with oxygenated ACSF + PiTx. O-LM hippocampal interneurons were identified by the location of their soma (*stratum oriens* of the CA1), their morphology (a spindle-shaped cell body horizontally oriented along the pyramidal layer) and their peculiar electrophysiological signature (the characteristic “sag” depolarizing potential in response to hyperpolarization currents injections and the typical “saw-tooth” shape of action-potential after-hyperpolarizations). Whole-cell patch-clamp recordings were obtained from CA1 O-LM interneurons with electrodes filled with an internal solution containing (in mM): K-gluconate, 120; KCl, 20; HEPES, 10; EGTA, 0; MgCl_2_6H_2_O, 2; Na_2_ATP, 2; spermine, 0.1. Stimulating pipettes filled with extracellular saline solution were placed in the stratum oriens to stimulate the axons of CA1 pyramidal cells.

Recordings were obtained using a Multiclamp 700B (Molecular Devices) amplifier and pClamp10.4 software. Data were sampled at 10 kHz, filtered at 3 kHz, and digitized by a Digidata 1440A (Molecular Devices). All data analyses were performed with custom written software in Igor Pro 6 (Wavemetrics). Values of membrane potential were corrected for liquid junction potential (~ −12 mV).

In control and test conditions the following parameters were measured in current clamp mode for 15 minutes before and 15 or 30 minutes after episode or 2 of 5 Hz stimulation. Apparent input resistance (R_in_) was tested by current injection (−120 pA; 800 ms); EPSPs were evoked at 0.1 Hz and the stimulus intensity (100 μs, 40–100 μA) was adjusted to evoke subthreshold EPSPs (4–10 mV). Synaptic transmission was measured between 2 and 4 ms from the onset of the EPSP. Short current step injections (+70/+120 pA, 100 ms) were applied at each sweep in order to measure the intrinsic excitability as the number of spikes over time. Series resistance was monitored throughout the recording and only experiments with stable resistance were kept (changes <10%). Before and after LTP induction, a protocol was designed in order to plot input-output curves by measuring action potentials number in response to incrementing steps of current pulses.

### Pharmacology

Excepted U73122 (1-[6-[[(17β)-3-Methoxyestra-1,3,5(10)-trien-17-yl]amino]hexyl]-1*H*-pyrrole-2,5-dione) that was added to intracellular solution, all drugs were bath applied. PiTx was purchased from Sigma-Aldrich; ZD-7288 [4-(N-ethyl-N-phenylamino)-1,2-dimethyl-6-(methylamino) pyrimidinium chloride], XE-991 dihydrochloride (10,10-*bis*(4-Pyridinylmethyl)-9(10*H*)-anthracenone dihydrochloride), NASPM trihydrochloride (*N*-[3-[[4-[(3-Aminopropyl)amino]butyl]amino]propyl]-1-naphthaleneacetamide trihydrochlo-ride), LY367385 (*S*)-(+)-α-Amino-4-carboxy-2-methylbenzeneacetic acid), U73122, chelerythrine chloride (1,2-Dimethoxy-12-methyl[1,3]benzodioxolo[5,6-*c*]phenanthridinium chloride) were obtained from Tocris and PIP2 (phosphatidylinositol-4,5-bisphosphate) from Sigma. ZD-7288 at 1 μM was applied at least 15 minutes before recording in order to obtain a full blockade of the HCN channel-mediated current (Gastrein et al., 2011).

### Voltage-clamp measurements of HCN and Kv7 channel-mediated current

HCN channel-mediated current was measured in voltage-clamp with voltage steps from −77 mV to −142 mV. The inward current was measured before and after 5 Hz stimulation. To measure the M-current mediated by Kv7 channels, voltage steps from −32 mV to −62 mV were applied (Lawrence et al., 2006), and the relaxing component of the M-current was then measured before and after 5 Hz stimulation.

### Synaptic and intrinsic plasticity protocol

LTP and LTP-IE was induced by stimulating the test synapse at 5 Hz during 10 minutes (i.e. 3000 stimulations). This protocol was repeated twice in most of the experiments. In some experiments, a 10 Hz stimulation during 5 minutes was also used (i.e. 3000 stimulations). Synaptic and intrinsic depotentiation was induced by negative pairing (Δt = −10 ms) between single postsynaptic spike and presynaptic stimulation delivered at a frequency of 10 Hz repeated in blocks of 6 stimulation (Incontro et al., 2021; Péterfi et al., 2012).

### Synaptic stimulation with θ firing patterns

θ oscillations were induced in acute slices by bath application of 25 μM carbachol. In these experiments, Ca^2+^ and Mg^2+^ concentrations were lowered to physiological values (i.e. 1.3 mM Ca^2+^ and 1 mM Mg^2+^), slices were kept intact (i.e. without surgical removal of area CA3) and recordings from CA1 pyramidal neurons were made in the presence of intact inhibition (without PiTx). Carbachol application induced spiking activity in the CA1 pyramidal neurons at the θ frequency during ~1 min. The action potential pattern was then collected and used to stimulate the CA1 to O-LM synaptic pathway via the ClampEx software.

### Morphological identification of biocytin-filled O-LM interneurons

A few O-LM interneurons were filled with biocytin to visualize the dendritic and axonal morphology of the recorded neurons. Biocytin (0.2-0.4%, Sigma Aldrich) was added to the pipette solution, and O-LM cells were filled at least for 20 min. Biocytin was revealed with streptavidin complex coupled to Alexa Fluor 488 (Thermo Fisher Scientific), and examined using confocal microscopy (Zeiss, LSM 780). The morphology of the neurons was reconstructed using ImageJ.

### Statistical analysis

EPSP slope, R_in_ and spike number were measured during the last 5 minutes of control or test periods. Pooled data are presented as mean ± SEM. Statistical comparisons were made using Wilcoxon or Mann-Whitney test as appropriate with Prism (GraphPad) software. Differences were considered as significant when p < 0.05 (*p < 0.05; **p < 0.01; ***p < 0.001).

## Author contributions

M.S., S.I. and D.D. designed research; M.S., Y.I., M.R. and S.I. performed research; M.S., N.A., S.I. and D.D. analyzed the data; M.S. and D.D. wrote the paper.

## Acknowledgements

Supported by INSERM, CNRS, AMU, Fondation pour la Recherche Médicale (DEQ20180839483 to D.D.), Agence Nationale de la Recherche (ANR-14-CE13-003 and ANR-21-CE16-0013 to D.D.). We thank Prof R. Nicoll and Prof D. Bowie for helpful comments on the manuscript.

---

**Supplementary Figure 1.**
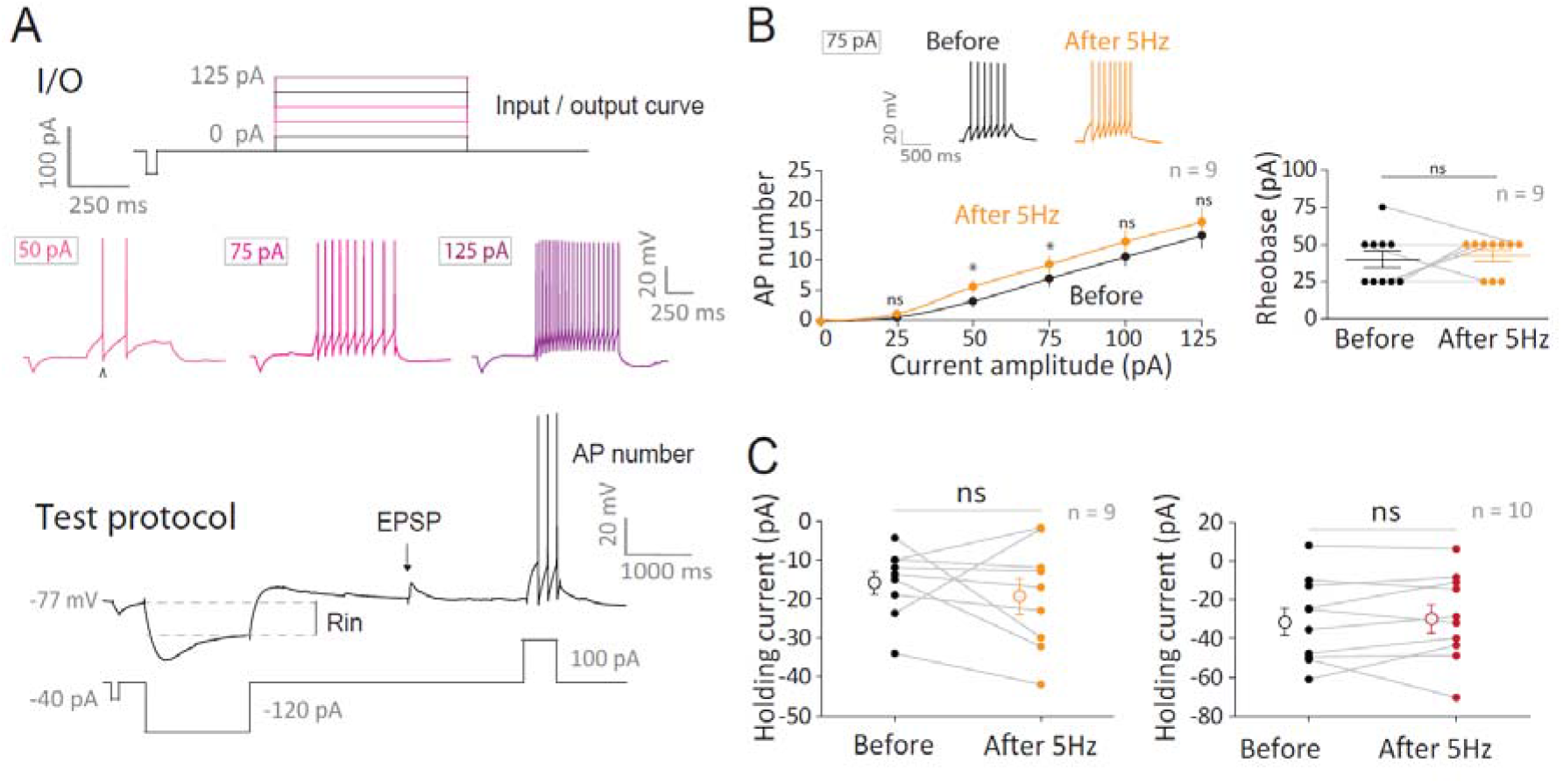
Protocols and intrinsic plasticity results. **A**. Current-clamp protocols. Top, input-output (I/O) curve: current steps (800 ms duration) from 0 to 125 pA were injected with increment of 25 pA. Bottom, test of synaptic and intrinsic changes. Input resistance (Rin) was measured with a long negative current pulse (−120 pA). EPSP was evoked by stimulation of pyramidal cell axon and the AP firing with a positive current pulse injection. **B**. Input-output curves (left) and rheobase (right) before and after a single 5 Hz episode. **C**. Holding current before and after a single (left) and two (right) episodes of 5 Hz stimulation. Wilcoxon and Mann-Whitney tests were used: ns, p>0.05; *, p<0.05; **, p<0.01.

**Supplementary Figure 2.**
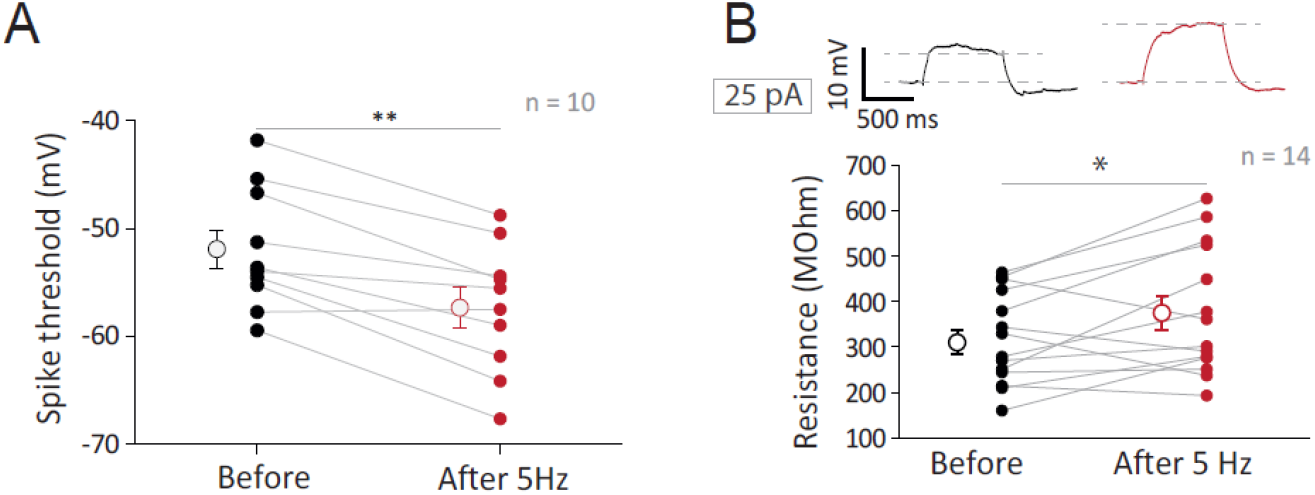
Change in AP threshold and in depolarizing voltage deflection. **A**. Hyperpolarization of the AP threshold after 2 episodes of 5 Hz stimulation. B. Stimulation at 5 Hz increases input resistance tested with depolarizing pulses of current. Top, traces; bottom, group data.Wilcoxon and Mann-Whitney tests were used: *, p<0.05; **, p<0.01.

**Supplementary Figure 3.**
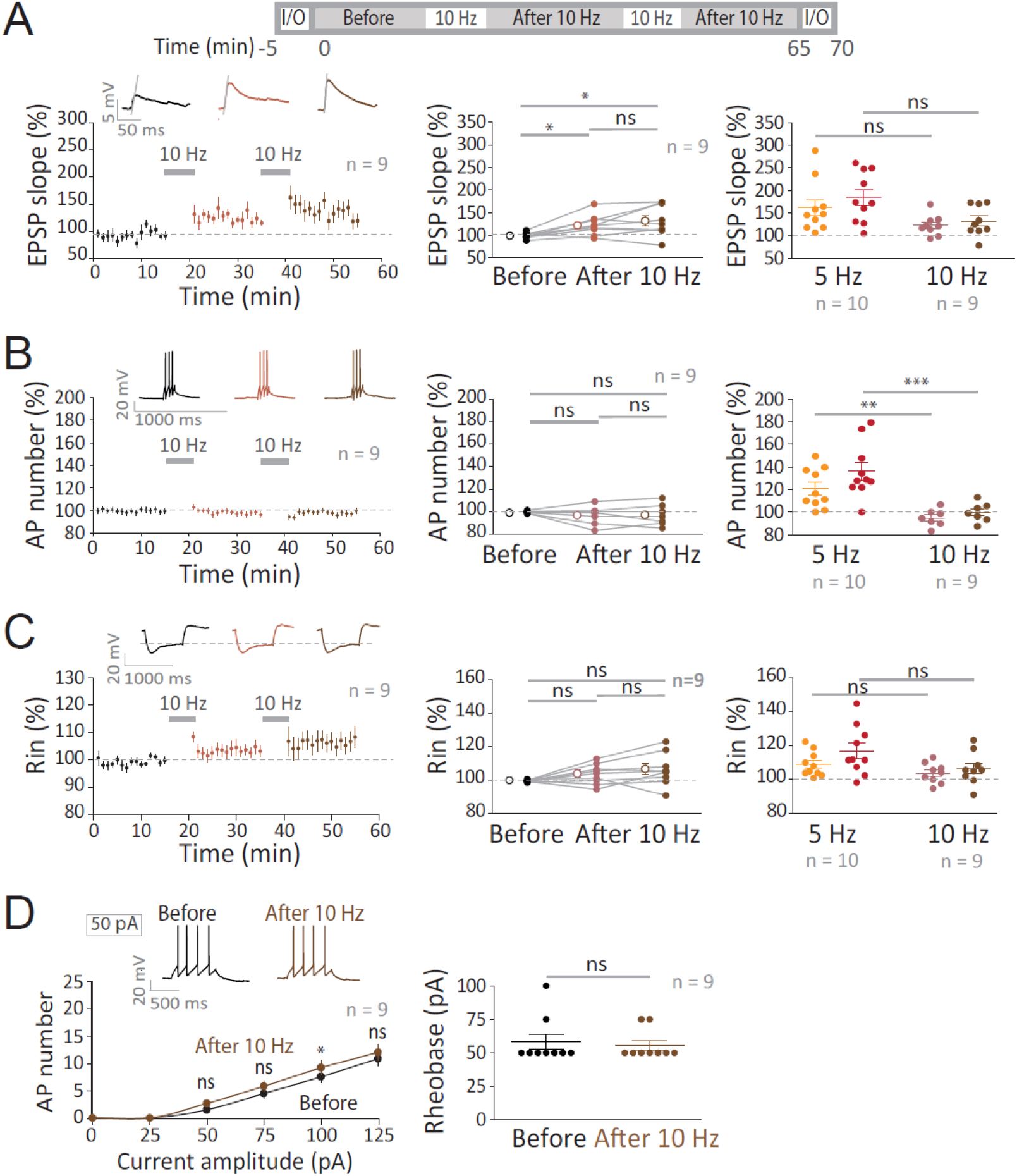
LTP-IE in O-LM interneurons is not induced by 10 Hz. **A**. Top, stimulation protocol. Bottom left, time-course of normalized EPSP slope before and after 10 Hz stimulation. Middle, group data and right, comparison with control. **B**. Left, time-course of normalized AP number before and after 10 Hz stimulation. Middle, group data and right, comparison with control. **C**. Left, time-course of normalized R_in_ before and after 10 Hz stimulation. Middle, group data and right, comparison with control. **D**. Input/output curves (left) and rheobase (right) before and after 10 Hz stimulation. Wilcoxon and Mann-Whitney tests were used: ns, p>0.05; *, p<0.05; **, p<0.01; ***, p<0.001.

**Supplementary Figure 4.**
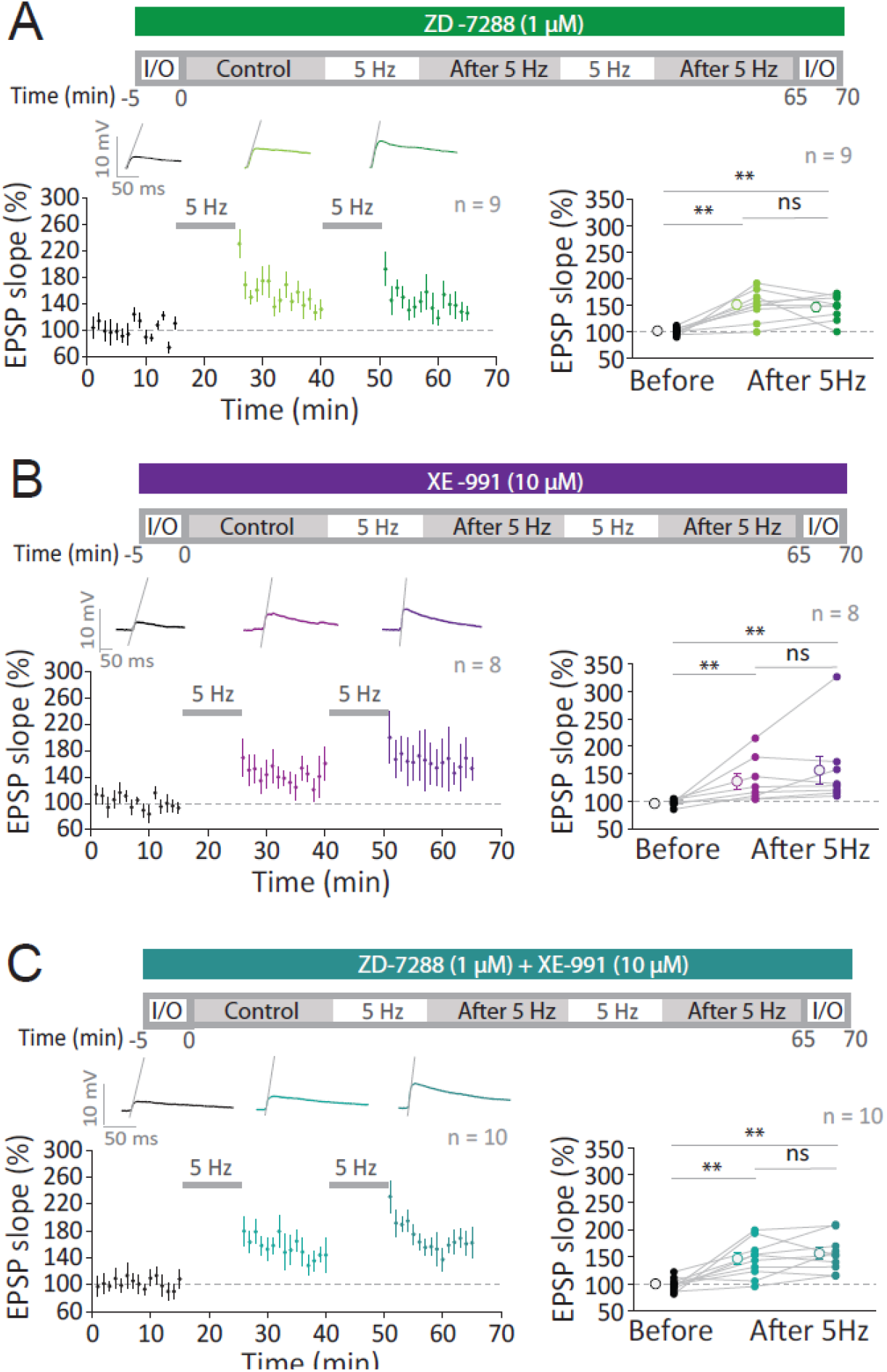
Normal LTP in the presence of HCN and Kv7 inhibitors. **A**. Top, stimulation protocol in the presence of ZD-7288. Bottom, time-course (left) and group data (right) of the EPSP slope in the presence of ZD-7288. **B**. Top, stimulation protocol in the presence of XE-991. Bottom, time-course (left) and group data (right) of the EPSP slope in the presence of XE-991. **C**. Top, stimulation protocol in the presence of ZD-7288 + XE-991. Bottom, time-course (left) and group data (right) of the EPSP slope in the presence of ZD-7288 + XE-991. Wilcoxon and Mann-Whitney tests were used: ns, p>0.05; **, p<0.01.

